# Fish presence alters amphibian and zooplankton communities in kettle lakes, but not hydrological connectivity

**DOI:** 10.64898/2026.01.22.700881

**Authors:** Ariane Barrette, Katrine Turgeon, Mariano J. Feldman, Guillaume Grosbois

## Abstract

Fishless lakes, critical drivers of biodiversity across freshwater landscapes, are becoming increasingly rare due to fish introductions. Although the impacts of fish introduction are well understood in high-elevation fishless lakes, their effects on fishless kettle lakes remain poorly understood. Many kettle lakes are disconnected from the surface water network and are therefore fishless. In this study, we examined how amphibian and zooplankton communities differ between fishless and fish-bearing kettle lakes by comparing 36 lakes in Québec, Canada. Some kettle lakes are hydrologically connected to surrounding aquatic ecosystems, allowing natural colonization by fish. We therefore also evaluated how amphibian and zooplankton communities differ between connected and disconnected kettle lakes. Fish presence was associated with differences at each stage of the amphibian life cycle. Reproductive calls of adult amphibians were detected regardless of fish presence, indicating that reproduction occurred in all lake types. However, the presence of fish was associated with fewer amphibian egg masses and lower larval abundance, and the absence of salamanders at the larval stage. Small-bodied zooplankton were more abundant in fish-bearing lakes, while overall species richness was lower. In particular, *Chaoborus americanus*, a large top-predatory zooplankton species, was found exclusively in fishless lakes. In contrast to fish presence, hydrological connectivity had no significant effect on most communities, except for adult American toads, adult wood frogs, and mink frogs’ larvae, which responded positively to the interaction between fish presence and connectivity. Based on our results, we recommend avoiding fish stocking of kettle lakes to preserve essential reproductive habitats for amphibians, maintain refuges for sensitive zooplankton species, and safeguard the spatial heterogeneity that underpins landscape-scale biodiversity.

## Introduction

Fishless lakes are critical drivers of biodiversity across freshwater landscapes (Stoks & McPeek, 2003). In the absence of fish, top-predator roles are filled by alternative taxa such as waterbirds, dragonflies, amphibians, benthic macroinvertebrates, and large zooplankton (Bradford et al., 1998; Hasan et al., 2023). However, fishless lakes are declining globally due to anthropogenic interventions, primarily fish introductions, that threaten their unique biodiversity and ecological functions (Miró & Ventura, 2013; Schilling et al., 2008).

Fish introductions into naturally fishless lakes can trigger cascading ecological changes. Whether through legal stocking or illegal release of sportfish or baitfish, these introductions can drastically restructure food webs, reduce biodiversity, and alter ecosystem functions (Sienkiewicz & Gąsiorowski, 2016; Tiberti et al., 2014). As efficient visual predators, introduced fish exerts strong top-down control on lower trophic levels (Tiberti et al., 2014). They prey preferentially on the most visible species, particularly those that are large, mobile and pigmented (Bradford et al., 1998; Hylander et al., 2012). Fish introductions into fishless lakes can therefore reduce the abundance and diversity of key freshwater taxa, such as amphibians, benthic macroinvertebrates, and large zooplankton, and even lead to local extinctions (Bradford et al., 1998; Tiberti et al., 2014). Furthermore, these introductions can affect other trophic levels, for example by increasing primary productivity if large grazing zooplankton are heavily preyed upon (Parker & Schindler, 2006; Vanni & Layne, 1997). These impacts have been primarily studied in high-elevation lakes, where fish are often absent due to limited natural colonization.

Newly introduced fish particularly affect alternative predators in naturally fishless lakes, including amphibians and large zooplankton, through direct predation or competition. For example, in high-elevation fishless lakes, fish introductions have caused major declines of the mountain yellow-legged frog (*Rana muscosa*) in the Sierra Nevada and in the long-toed salamander (*Ambystoma macrodactylum)* in northern California through predation (Knapp et al., 2001; Welsh Jr et al., 2006). However, other factors like amphibian size, palatability and anti-predatory behaviour can mitigate the impacts of these introductions (Hecnar & M’Closkey, 1997; Kats et al., 1988; Welsh Jr et al., 2006). As for zooplankton, introduced fish in fishless lakes exert selective predatory pressure on larger and more visible zooplankton, such as *Chaoborus* species, large-bodied cladocerans, and calanoid copepods (Bradford et al., 1998; Holmes et al., 2017). As a result of reduced interspecific competition for food resources with large-bodied zooplankton, small-bodied taxa, such as smaller cladocerans, cyclopoid copepods, and rotifers, can increase in abundance (Knapp et al., 2001; Tiberti et al., 2014).

In addition to anthropogenic fish introductions, hydrological connectivity allows the natural colonization of lakes by fish. According to metacommunity theories, connectivity promotes the dispersal of aquatic organisms across connected lakes, thereby supporting ecosystem resilience and species recolonization following disturbances (Bouvier et al., 2009; Verboom et al., 1993). In contrast, aquatic species dispersal is strongly limited in disconnected lakes, which remain isolated from the regional metacommunity.

Consequently, disturbances in these ecosystems can have a greater impact on local communities (Haddad et al., 2015). Nevertheless, isolation also creates unique ecological conditions that may foster the emergence of endemic species (Fuke et al., 2024; Nazarov et al., 2023; Power et al., 2024). Thus, disconnected lakes can represent important biodiversity hotspots and may enhance biological productivity (Blackburn-Desbiens et al., 2023).

Kettle lakes on eskers originate from fluvioglacial formations characteristic of northern regions: the eskers and moraines. As glaciers retreated during the last glaciation, blocks of ice broke off and were buried within the sandy deposits of these formations, creating depressions that later evolved into kettle lakes (Veillette et al., 2004). Many of these lakes are disconnected from the surface water network and are fed only by groundwater and precipitation, which prevents natural colonization by fish (Schilling et al., 2008; Veillette et al., 2004). Other kettle lakes are hydrologically connected to surface or groundwater networks and can support indigenous fish populations (Veillette et al., 2004). In addition, several lakes have been stocked with fish, legally or illegally, with consequences that remain poorly understood. The biodiversity and ecology of kettle lakes are understudied, with only a few studies focusing on waterbird and macroinvertebrate populations (Grosbois et al., 2025; Hasan et al., 2023; Schilling et al., 2009a). Previous studies have shown that fishless kettle lakes provide habitat that favors certain waterbird species, such as the common goldeneye (*Bucephala clangula*) and the Canada goose (*Branta canadensis*). However, fish stocking in these lakes has been found to reduce macroinvertebrate diversity and abundance (Grosbois et al., 2025; Hasan et al., 2023; Schilling et al., 2009a). Despite these findings, information remains lacking on the impact of fish on apex predator taxa in originally fishless lakes, particularly amphibians and zooplankton.

Our study evaluated the abundance, diversity and species composition of amphibians and zooplankton between fishless and fish-bearing kettle lakes. We hypothesized that amphibians would be less abundant and less diverse in fish-bearing lakes and that species composition would differ between fishless and fish-bearing lakes. We also hypothesized that larger predatory zooplankton would be less abundant in fish-bearing lakes, whereas smaller zooplankton would be more abundant due to top-down control by fish. We expected fish presence to reduce zooplankton diversity, with differences in species composition between fishless and fish-bearing lakes. Because kettle lakes can be either connected or isolated from the surrounding surface water network, conditions that influence amphibian and zooplankton dispersal across aquatic ecosystems, we also evaluated how connectivity affects their diversity, abundance, and species composition. We hypothesized that connectivity would mitigate the impact of fish presence on these communities by facilitating the exchange of individuals.

## Methods

### Study sites

The study was conducted from May to August 2024 in the Abitibi-Témiscamingue region of Québec, Canada, an area where eskers are abundant (Fig. 1) (Veillette et al., 2004). The study area is located between 48°23’ and 48°58’ and between 77°24’ and 78°40’ (WGS84) within the balsam fir (*Abies balsamea* L. Mill.) – white birch (*Betula papyrifera* Marsh.) bioclimatic domain (MFFP, 2021). The climate is humid continental, with a mean annual temperature of 1.5 °C and an average annual precipitation of 923.2 mm for the 1981–2010 period (MELCCFP, 2025; Natural Resources Canada, 2022). The study was carried out on the Saint-Mathieu - Berry, Vaudray-Joannès, Launay and Lac Despinassy eskers, as well as on the Harricana moraine; given their similar geological composition, we refer to both landforms collectively as “eskers”. Numerous kettle lakes are located on these eskers and are subjected to intense anthropogenic pressures such as forest harvesting, mining and recreational activities (Guimond et al., 2024), reflecting complex interactions with their watersheds (Grosbois et al., 2023). These dimictic lakes are ice-covered from late November to early May on average and undergo two turnovers, in spring and autumn (Brönmark & Hansson, 2018; Grosbois et al., 2024).

**Figure 1.**
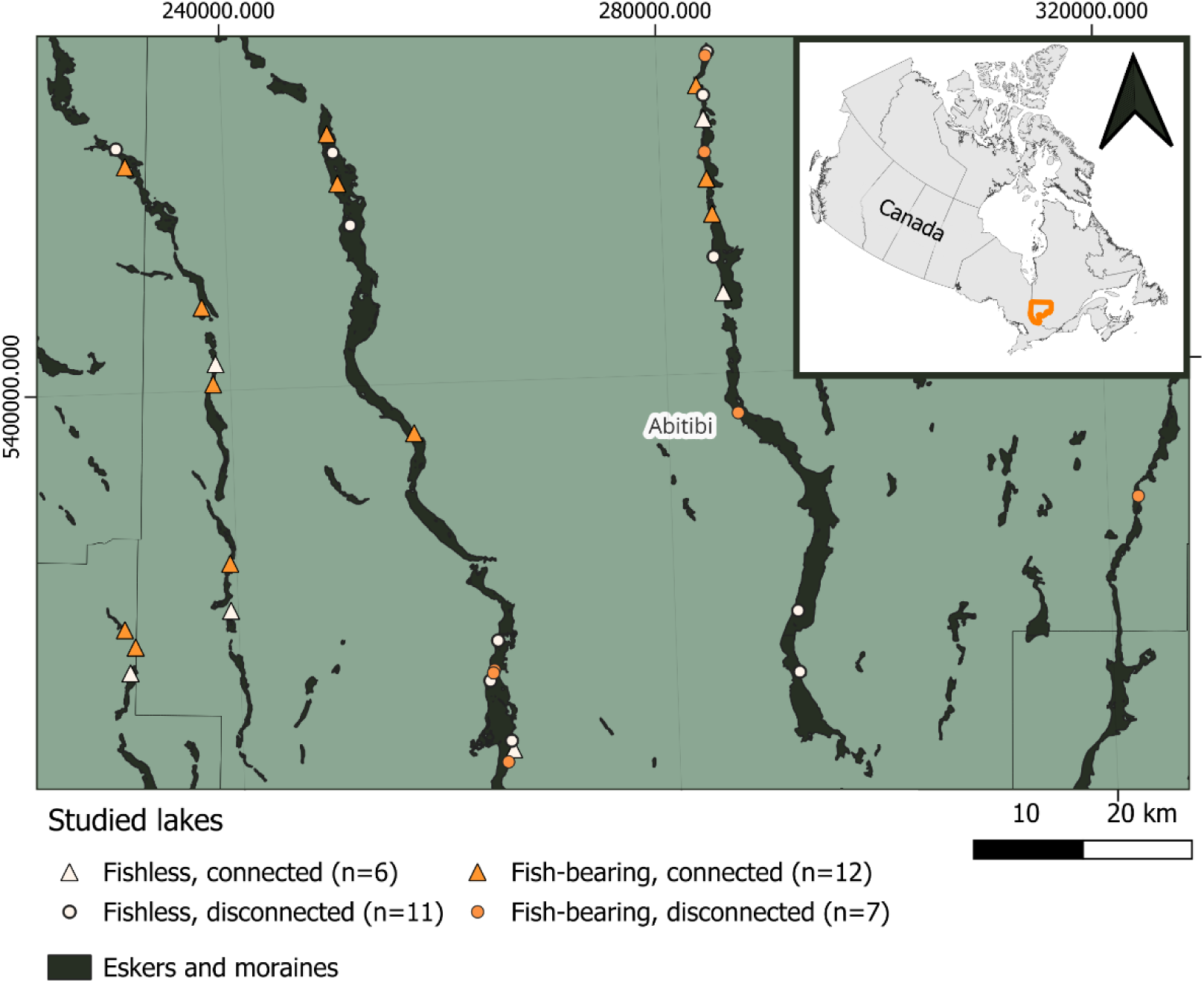
Location of the 36 studied lakes

### Study design

We sampled 36 kettle lakes located on eskers, including 17 fishless lakes and 19 fish-bearing (Fig. 1). Among the fishless lakes, six were connected to the surface water network and 11 were disconnected. Of the fish-bearing lakes, 12 were connected to the surface water network, and seven were disconnected from it. Lakes were selected based on documented fish populations (Hasan et al., 2023), connectivity to the surface water network, surface area size (<7 ha) and accessibility (e.g., distance to the nearest road). To minimize spatial dependence, lakes were separated by at least 1 km. Fish presence or absence was confirmed through a field survey conducted in this study. A lake was classified as fish-bearing if at least one fish was captured, and considered connected if a permanent stream linked it to another connected lake or to a wetland that itself drained into the surface water network. Connectivity to the surface water network was assessed using geospatial datasets, including CMHPQ (MELCCFP, 2023) and hydrographic derivatives from LiDAR.

### Physico-chemical sampling

We measured key limnological variables expected to influence amphibian, fish, and zooplankton assemblages (Dodd, 2023; Marium et al., 2023; Yousef et al., 2024). We collected physico-chemical data in the littoral zone of each sampled lake at one sample site. Specific conductivity (µS/cm), pH, dissolved oxygen concentration (µmg/L), and dissolved oxygen saturation (%) were measured at 30 cm depth using a multiparameter probe (RBR Concerto, Ottawa, Canada). The euphotic zone was characterized by measuring light attenuation in water using an underwater light sensor and an air control light sensor (LI-1500, LiCor, Lincoln, USA), with measurements taken at the water surface and at three additional depths, incrementing by 10 cm. Suspended algal biomass in lake water was estimated from chlorophyll a (chl-a) concentration. To measure chl-a, 1 L of epilimnion water was collected from each lake in a brown opaque Nalgene bottle and filtered in triplicate through glass fiber filters (GF/F, 47 mm, 0.7 µm) at the Groupe de Recherche en Écologie de la MRC-Abitibi (GREMA) laboratory of the Université du Québec en Abitibi-Témiscamingue (UQAT). The filters were wrapped in aluminum foil to protect them from light and stored at -80°C until analysis. To measure total phosphorus (TP) and total nitrogen (TN), four 50 mL glass vials were acid-washed (10% HCl) for 24h, oven-dried at 200°C, and then filled with lake water. For dissolved organic carbon (DOC) and dissolved inorganic carbon (DIC), four 50mL brown vials were filled with lake water filtered through a 0.45 µm filter. These DOC-DIC vials were previously combusted at 550 °C for at least 4 hours to remove carbon. All water samples were collected at a depth of 30 cm. Nitrogen samples were analyzed with a continuous flow analyzer (OI Analytical Flow Solution 3100 ©, College Station, USA) using an alkaline persulfate digestion method, coupled with a cadmium reactor, following a standard protocol (Patton & Kryskalla, 2003). Phosphorus samples were analyzed using a standard protocol (Wetzel & Likens, 2000) and DOC-DIC samples with an OI Analytical Aurora 1030W TOC Analyzer (https://www.oico.com/1030W) (Yellow Springs, USA) using a persulfate oxidation method. All water samples were analyzed at the GRIL analytical laboratory at the Université du Québec à Montréal (UQAM).

### Biological data compilation

#### Fish surveys

Presence/absence of fish has been estimated in each lake by deploying three baited minnow traps (Gee-Feets G-40, hole entrance of 2.5 cm) and one fyke net (1.22 m high, 1.22 m wide, mesh size of 2 mm) overnight in each lake on one occasion between mid-May and late July 2024. We used both fishing gears because they sample different habitats and size classes of the fish community, thereby improving the detection of a broad range of species present (Fischer & Quist, 2014). We identified, measured, and weighted each captured individual before release.

#### Amphibian surveys

We used a variety of methods to assess the amphibian community in esker lakes, including acoustic recorders (for male adults), visual surveys (for egg masses), and captures with minnow traps and a fyke net (for larvae and newts). We conducted anuran call surveys based on the breeding phonologies of the species expected to occur within the study area, including wood frogs (*Lithobates sylvaticus*), mink frogs (*Lithobates septentrionalis*), green frogs (*Lithobates clamitans*), northern spring peepers (*Pseudacris crucifer crucifer*) and American toads (*Anaxyrus americanus)* (Rodrigue & Desroches, 2018). Calling surveys were conducted in two periods: at the beginning of May (starting April 29^th^) during the breeding season of early-breeding species (first sampling period), and 40 days later during the breeding season of later-breeding species, in June (starting June 7^th^; second sampling period) (Bouthillier et al., 2015; Feldman et al., 2023). We deployed one acoustic recorder (SM4 Song meter, Wildlife Acoustics, Maynard, USA) at each lake at 2 to 10 m from the lake shore, depending on the position of the first available tree and the most suitable unobstructed location for maximizing sound capture, and at a height of 1.5 m on a tree. Following Feldman et al. (2023), we programmed acoustic recorders to record for 3 minutes per hour from 21:00 to 02:00 for seven consecutive days for each sampling period (total of fourteen days of call survey per lake) (Annich et al., 2019; Bevier et al., 2004). Calls were then identified listening to audio sample by the same person (A. Barrette) to reduce observer bias. In total, 84 audio samples were listened per lake, representing 252 minutes of recordings per lake.

After listening to the full length of the 3-min recordings, we followed the North American Amphibian Monitoring Program (2005) to annotate the calling scores. Calling scores were assigned as follows: 0 if no individuals were heard; 1 if individuals can be counted with no overlap in calls; 2 if a few individuals can be counted despite overlapping calls; and 3 if a chorus with individuals can not be counted. We retained the maximum calling score observed for each species in each lake on each night. These call indices were then considered as the abundance of each species.

At the end of each sampling period, after the breeding season of either early or later-breeding species, we conducted visual surveys to quantify the relative abundance of amphibian egg masses (Campbell Grant et al., 2005; Egan & Paton, 2004; Heyer, 1994). Visual surveys were conducted between 08:00 and 17:00 at the end of each sampling period (ie., May 13^th^ to 17^th^ and June 24^th^ to 28^th^). Two observers, equipped with polarized sunglasses to reduce surface glare and improve visibility in the water, surveyed a 200 m shoreline transect in each lake, walking in opposite directions from a common starting point. Each observer independently identified and counted egg masses along a shoreline strip within ≥2 m wide, extending the shoreline strip when necessary to reach a water depth of 0.5m.

To characterize amphibian larvae communities, we quantified the relative abundance of tadpoles and salamander larvae (*Ambystoma* sp.). Larvae were sampled simultaneously as fish by deploying three baited minnow-traps overnight equipped with glow sticks and one non-baited fyke net in each lake. These fishing gears also permitted the capture of adult eastern newts (*Notophthalmus viridescens viridescens*). Glow sticks were used to improve capture success, as larval amphibians are attracted to light (Bennett et al., 2012). All individuals were identified to species following Rodrigue and Desroches (2018), except salamanders, which were identified to genus, then measured, weighed, and counted before being released.

Finally, we calculated the amphibian species richness and the Jaccard index as biodiversity indices for each lake. Any species identified at any life stage during sampling was considered present for species richness and Jaccard index calculations. Jaccard index was calculated using the R package vegan version 2.6-10 (Oksanen et al., 2025).

#### Zooplankton survey

The diversity and relative abundance of zooplankton in each lake were estimated with three vertical hauls using a 50 µm mesh zooplankton net of 25 cm diameter in triplicate at the deepest point of the lake. All lakes were visited within seven days at the beginning of August. Zooplankton samples were preserved in ethanol (final concentration 70%) in 125 ml Nalgene bottles. If necessary, samples were subsampled with a Folsom plankton divider to obtain approximately 400 individuals, excluding nauplii, female calanoids, male cyclopoids, and juvenile copepods. We then identified zooplankton to the lowest taxonomic level possible and counted all individuals using a binocular (Zeiss, Discovery V12, Oberkochen, Germany) in the GREMA-UQAT laboratory (Haney et al., 2013). Mean relative abundance was calculated from the three replicates and reported per 1000 L of water filtered. Individuals identified to the species were selected to calculate lake-species richness, Shannon-Weiner and Simpson’s indices, Pielou’s evenness, as well as Bray-Curtis dissimilarity, with R package vegan.

### Data analysis

#### Physico-chemistry

DIC and dissolved oxygen saturation data were removed from the analysis as they were highly correlated with other variables (|r| > 0.7; See Fig. S1 in *Supplementary material*). Other physico-chemical variables were compared between lake types using either a Two-Sample t-test, a Welch Two-Sample t-test, or a Wilcoxon’s test, depending on the data’s homoscedasticity and normality. Data were log-transformed (log_10_(x+1)) to achieve normality and homoscedasticity if necessary.

#### Amphibian communities

To assess how fish presence, hydrological connectivity, and their interaction were associated with amphibian reproduction, we modelled egg mass abundance and larvae abundance (response variables) using zero-inflated negative binomial models (R package *glmmTMB*, v. 1.1.11, M. Brooks et al., 2017), fitting separate models for each species. Fish presence (i.e., fishless; fish-bearing), connectivity (i.e., connected; disconnected), their interaction and scaled physico-chemical variables (i.e., mean of 0, SD of 1) were included in models and backward elimination was used to obtain the most parsimonious model (See Table S1 in *Supplementary material*). The exact length of the transect for the egg mass survey or the fishing hours was accounted for using an offset in the formula. Physico-chemical variables included in the models were previously preselected based on their biological relevance, results from the Boruta feature selection algorithm, and Akaike Information Criterion (AIC) values (See Table S1 in *Supplementary material*). Boruta was conducted using R packages Boruta (v. 8.0.0, Kursa & Rudnicki, 2009) and randomForest (v. 4.7-1.2, Breiman et al., 2024), and AIC using MuMIn (v. 1.48.4, Bartoń, 2010).

The relative abundance of egg masses was converted to presence/absence data because models could not be fitted successfully using the combined relative abundance across all amphibian species. To do so, if at least one observer identified an egg mass of a species, the species was considered to be present. We conducted a Generalized Linear Model (GLM) with a binomial family to assess the effects of fish presence, connectivity, their interaction, species identity and preselected physico-chemical variables on the probability of egg mass presence. The exact length of the transect was considered using an offset in the formula. Backward elimination was then applied to derive the most parsimonious model. To verify assumptions, the mod.check function from the R package DHARMa (v. 0.4.7, Hartig et al., 2024) was used.

A linear model was also conducted to assess the effects of fish presence, connectivity, their interaction, and preselected physico-chemical variables on total relative larval abundance. We then used backward elimination to obtain the most parsimonious model. Relative larval abundance was log-transformed to achieve the assumptions of linear models.

Generalized Additive Models (GAMs) were conducted using the GAM ordered categorical family (OCAT) to assess the effects of fish and connectivity and their interaction on the calling score of each anuran species. Sampling periods were treated independently. Fish presence, connectivity and their interaction were set as linear predictors, Julian day as a smooth (non-linear) predictor and the basis dimension *k* was set to 10. The R package mgcv (v. 1.9-1, Wood, 2000) was used and the functions k.check and gam.check were used for model checking. Calling scores were previously summed to 1, as GAMs require positive values.

Finally, Wilcoxon tests were made to compare amphibian species richness between fishless and fish-bearing lakes, and to compare amphibian and fish species richness between connected and disconnected lakes. The Jaccard index matrix was plotted using non-metric multidimensional scaling (NMDS), and a PERMANOVA was conducted using the adonis function in the vegan R package to assess the effects of fish presence and connectivity. Two lakes were excluded from the amphibian diversity analysis because the recording devices had been stolen.

#### Zooplankton communities

Two-sample t-test or Wilcoxon’s test were performed to compare zooplankton family abundances, total zooplankton relative abundances, Pielou’s evenness, and Shannon-Weaver and Simpson’s indices between lake types (fishless vs fish-bearing lakes, connected vs disconnected lakes), depending on homoscedasticity and normality of the data. Data were log-transformed when necessary to meet model assumptions. A GLM with a binomial family was also conducted on the presence of *Chaoborus*, with fish presence, connectivity and their interaction included as independent variables, and backward elimination was used to obtain the most parsimonious model.

The effects of fish presence, connectivity and their interaction on species richness were analysed using a linear model and backward elimination. The Bray-Curtis dissimilarity matrix was plotted using NMDS and a PERMANOVA was conducted to compare species assemblages among lake types. All statistical analyses were completed using R 4.3.3 (R Core Team, 2024) and considered to be significant if p < 0.05.

## Results

### Lake characteristics

Lakes were similar in their physico-chemistry, except for pH and specific conductivity (Table 1). pH was one unit higher in fish-bearing lakes (p = 0.03, Table 1). Specific conductivity was about five times higher and more variable in fish-bearing lakes compared to fishless lakes (t = -3.711, p = 0.001, Table 1). Physico-chemical characteristics were also similar between connected and disconnected lakes, but specific conductivity was higher in connected lakes (p = 0.007; Table 1).

**Table 1.** Physico-chemical characteristics of the 36 studied lakes (mean ± standard deviation [min,max]). p-values were obtained from a Two Sample t-test (normally distributed data with equal variances), a Welch Two Sample t-test (normally distributed data with unequal variances) or a Wilcoxon test (non-normally distributed data) between fishless and fish-bearing lakes, and between disconnected and connected lakes. If needed, data were log-transformed before applying the statistical test. Significant differences between lake types are highlighted in bold and with an asterisk (*).

| Lake physico-chemistry | Lake type |  |  |  |  |  |
| --- | --- | --- | --- | --- | --- | --- |
|  | Fishless (n=17) | Fish-bearing (n=19) | p-value | Disconnected (n=18) | Connected (n=18) | p-value |
| Dissolved oxygen concentration ($\mu\text{mol/L}$) | 236.8 ± 22.9<br>[196.8 : 270.2] | 241.4 ± 23.3<br>[196.8 : 292.4] | 0.58 | 233.2 ± 16.4<br>[206.5 : 260.6] | 245.8 ± 27.3<br>[196.7 : 292.4] | 0.11 |
| pH | 6.5 ± 0.8<br>[5.7 : 8.0] | 7.4 ± 1.3<br>[5.6 : 9.9] | **0.025\*** | 6.7 ± 0.8<br>[5.7 : 8.8] | 7.8 ± 1.4<br>[5.6 : 9.9] | 0.18 |
| Specific conductivity ($\mu\text{S/cm}$) | 6.2 ± 3.6<br>[2.3 : 13.4] | 33.6 ± 37.3<br>[2.8 : 113.2] | **0.001\*** | 13.2 ± 23.2<br>[2.3 : 97.3] | 29.7 ± 35.6<br>[3.6 : 113.2] | **0.007\*** |
| Dissolved organic carbon (mg/L) | 7.4 ± 3.4<br>[3.0 : 14.0] | 6.8 ± 4.2<br>[1.1 : 15.9] | 0.63 | 6.4 ± 3.3<br>[2.7 : 14.0] | 7.7 ± 4.2<br>[1.1 : 15.9] | 0.36 |
| Chlorophyll-a<br>(µg/L) | 6.3 ± 4.1<br>[1.7 : 14.9] | 5.5 ± 4.8<br>[1.8 : 23.1] | 0.38 | 5.5 ± 3.5<br>[1.7 : 14.9] | 6.3 ± 5.3<br>[1.8 : 23.1] | 0.79 |
| Total phosphorus<br>(µg/L) | 21.4 ± 9.5<br>[8.5 : 42.2] | 18.0 ± 7.3<br>[6.7 : 39.7] | 0.32 | 18.4 ± 9.0<br>[8.5 : 42.2] | 20.8 ± 7.9<br>[6.7 : 39.7] | 0.20 |
| Total nitrogen<br>(ppm) | 0.4 ± 0.1<br>[0.2 : 0.8] | 0.4 ± 0.2<br>[0.1 : 1.0] | 0.62 | 0.4 ± 0.1<br>[0.2 : 0.8] | 0.4 ± 0.2<br>[0.1 : 1.0] | 0.24 |
| Photic zone<br>(m) | 3.2 ± 2.0<br>[1.1 : 9.4] | 3.8 ± 1.9<br>[1.2 : 7.4] | 0.30 | 3.8 ± 1.5<br>[1.7 : 6.7] | 3.2 ± 2.3<br>[1.1 : 9.4] | 0.079 |

### Fish community

Across the 19 fish-bearing lakes, ten native fish species were caught. The most common species was brook stickleback (*Culaea inconstans*), followed by brown bullhead (*Ameiurus nebulosus*), northern redbelly dace (*Chrosomus eos*), and yellow perch (*Perca flavescens*), and we also captured occasional species of the Cyprinidae, Cottidae, Salmonidae, and Catostomidae families in some lakes (See Table S2 in *Supplementary material*). We captured ten species in connected lakes (mean ± SD, 2.2 ± 1.3 per lake; Table S2) and six species in disconnected lakes (1.7 ± 1.3 per lake), but this difference was not significant (p = 0.2).

### Amphibian communities

#### Egg masses

Egg masses of two early-breeders’ species, the spotted salamander (*A. maculatum*) and the wood frog, were identified and counted, as well as one late-breeders’ species, the mink frog. The presence of fish significantly affected both the presence of egg masses (Fig. 2a) and the abundance of egg masses of early-breeders but not of late-breeders. The model showed that spotted salamander egg masses were significantly less abundant in fish-bearing lakes (estimate = -2.55, p = 0.027; Table S3). Wood frog egg masses were also less abundant in fish-bearing lakes, but the effect only approached statistical significance (estimate = -3.95, p = 0.06; Table S3). During the second sampling period in June, only a few egg masses of the mink frog, a late-breeder, were recorded. Fish presence was not correlated with mink frog egg masses’ abundance (estimate = -0.14, p = 0.82; Table S3), nor was connectivity (estimate = -0.10, p = 0.87; Table S3).

**Figure 2.**
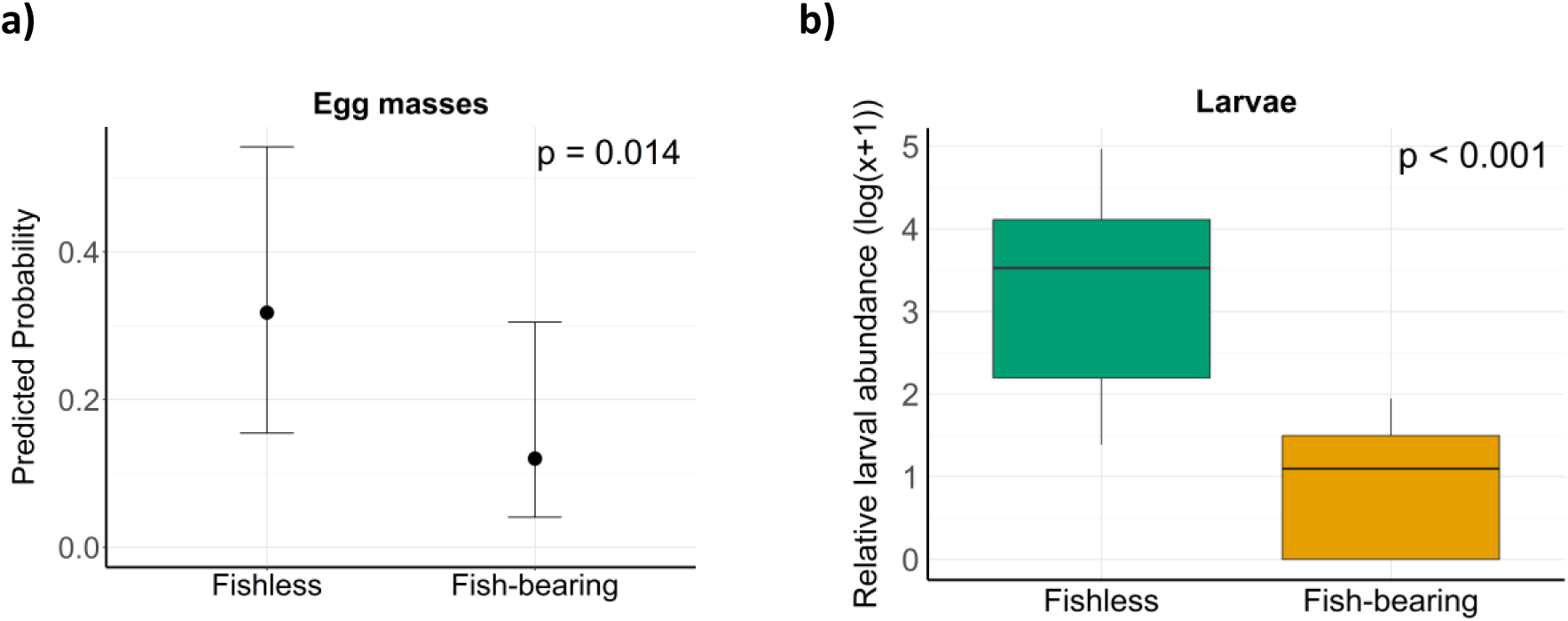
Impacts of fish presence on the life cycle of amphibians. a) Predicted probabilities of egg mass presence in fishless and fish-bearing lakes, with 95% confidence intervals, when fish presence, connectivity and amphibian species are included in the model. All amphibian species were combined. b) Boxplot of larval abundance (all species combined) captured over a 24-hour period in fishless and fish-bearing lakes. Abundances have been rounded upwards before being log(x+1)-transformed.

The GLM showed that the presence of fish reduced the odds of observing an egg mass by 71%, all species combined (log-odds = -1.23, odds = 0.29, p = 0.014; Table S3). Connectivity did not have a significant effect on the presence of egg masses (log-odds = 0.46, odds = 1.59, p = 0.34; Table S3). The predicted probability of observing an egg mass (presence) in fishless lakes is 32%, compared with 12% in fish-bearing lakes, when fish presence, connectivity, and amphibian species are included in the model (Fig. 2a).

#### Amphibian larvae

Amphibian larvae were more abundant in fishless lakes than in fish-bearing lakes when all species are combined (Fig. 2b), and the difference was more pronounced in disconnected lakes. Overall, we captured 725 larvae across fishless lakes and 120 in fish-bearing lakes. Mink frog tadpoles were the most frequently captured larvae (recorded in 18 lakes) and represented one-third of the catch. Salamanders, wood frogs, green frogs, and northern spring peepers larvae and adult eastern newts were also captured.

Fish presence was strongly associated to the decrease of larval abundance when all species were combined (catch per 24-hour, log-transformed; estimate = -2.11, p < 0.001, Fig. 2b; Table S3). Fish presence also significantly decreases mink frog tadpole relative abundance by 98% in disconnected lakes (estimate = -3.69, p < 0.001; Table S3) but by 56% in connected lakes (estimate = 2.89, p = 0.031; Table S3). No significant effect of fish presence, nor connectivity, was found to explain green frog larvae abundance, but this species was only captured in six lakes, two of which were fishless. Salamander, wood frog, green frog, and northern spring peeper larvae were too rare to be analyzed statistically, but salamander larvae were only found in fishless lakes.

DOC was found to have a small, but positive impact on total relative larval abundance when all species were pooled (estimate = 0.51, p = 0.006; Table S3). Fish presence and dissolved organic carbon accounted for 60% of the variance in log-transformed relative larval abundance.

#### Male adults

The influence of fish presence or connectivity on adult male anuran calling scores differed across species. During the first sampling period, calls of three early-breeders were detected: the northern spring peeper, the wood frog, and the American toad. The northern spring peeper was calling in all the sampled lakes, while the wood frog and American toad called in 97% and 56% of the sampled lakes respectively. During the second sampling period, northern spring peeper was heard in 94% of the sampled lakes and American toad in 41%. Two late-breeders were also heard on the recordings: the mink frog in 88% of the sampled lakes and the green frog in 32%.

The American toad calling scores were higher in fish-bearing lakes during the first sampling period (estimate = 2.04, p = 0.002; Table S3), but not during the second (estimate = -1.66, p = 0.016; Table S3). Mink frog calling scores were also higher in fish-bearing lakes (estimate = 0.72, p = 0.045; Table S3). Wood frog (estimate = -1.35, p < 0.001; Table S3) and green frog (estimate = -1.51, p = 0.006; Table S3) calling scores were lower in fish-bearing lakes, while no clear pattern was observed for the northern spring peeper.

There was also no clear pattern of the impact of connectivity on calling scores of most species. However, connectivity had a positive impact on the American toad calling scores during the first sampling period (estimate = 1.58, p = 0.019; Table S3). During the second sampling period, the interaction between fish presence and connectivity significantly affected American toad and wood frog calling scores (respectively: estimate = 3.19, p < 0.001; estimate = 1.157, p = 0.03; Table S3).

#### Diversity and composition

We recorded a total of seven amphibian species during the sampling period (mating calls of adult males recorded at night and active sampling of egg masses and amphibian larvae). These species included *Ambystoma* sp, wood frog, mink frog, green frog, northern spring peeper, American toad and eastern newt.

Amphibian species richness did not differ (p = 0.5) between fishless and fish-bearing lakes, with a mean species richness of 4.4 ± 0.9 and 4.2 ± 1.1 species/lake, respectively. Species richness was also not different between connected and disconnected lakes. Overall, fish presence and connectivity had a marginally non-significant effect on amphibian community composition, as measured by the Jaccard index (Fish: F_(1, 31)_ = 2.30, p = 0.087; Connectivity: F_(1, 31)_ = 2.48, p = 0.065) (Fig. 3).

**Figure 3.**
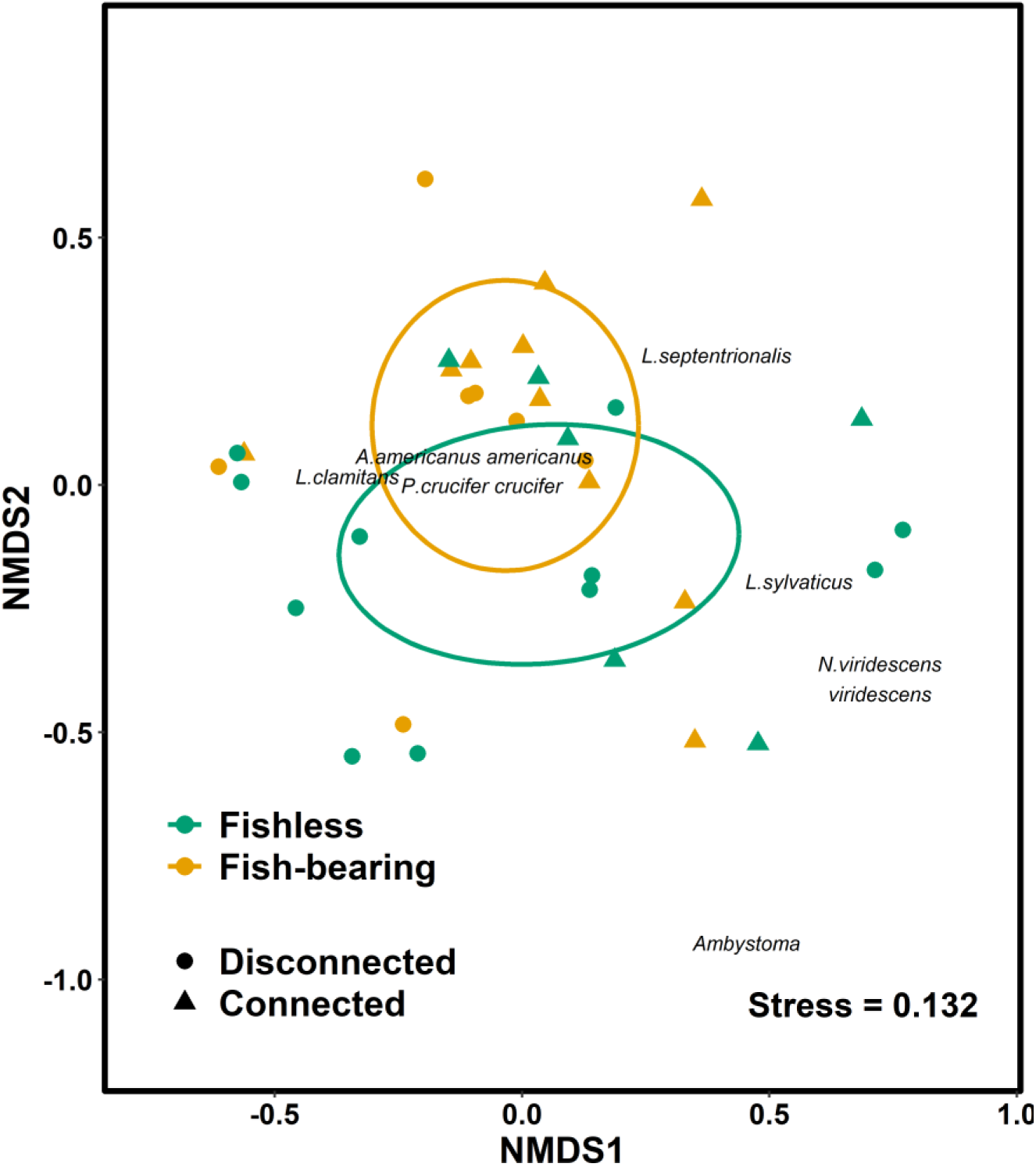
Non-metric multidimensional scaling (NMDS) plot based on the Jaccard index for amphibian communities in summer 2024. Green: Fishless lakes (n=17), orange: Fish-bearing lakes (n=17), circle: Disconnected lakes (n=18), triangle: Connected lakes (n=16). Points were jittered slightly to reduce overlap and improve visibility.

### Zooplankton communities

#### Diversity, composition, abundance

Zooplankton abundance was generally low in all sampled lakes (mean = 4.5 ± 7.3 individuals/L, range = 0.08 to 40.9 individuals/L) and did not differ between fishless and fish-bearing lakes (t=-1.343, p = 0.19). The zooplankton communities were composed of 14 species and three species of *Chaoborus* (Diptera) (See Table S4 in *Supplementary material*). Two different genera of cyclopoids were identified across the 36 lakes sampled, five calanoids and seven cladocerans. *Bosmina longirostris, Diaphanosoma brachyurum, Holopedium gibberum* and female calanoids were the four most encountered taxa.

*Chaoborus* larvae were less likely to be captured in fish-bearing lakes (log-odds = -2.20, odds = 0.11, p = 0.005; Table S3). Indeed, *Chaoborus americanus* was observed only in fishless lakes. Other *Chaoborus* species were also observed in fish-bearing lakes. Furthermore, Bosminidae was significantly more abundant in fish-bearing lakes (Unilateral Wilcoxon test, p = 0.03) as well as Diaptomidae (Unilateral Wilcoxon test, p = 0.05). No patterns were detected for other zooplankton families.

Zooplankton species richness was greater in fishless lakes than in fish-bearing lakes (log-transformed species richness, estimate = -0.22, p = 0.037; Table S3). Mean species richness in fishless lakes was 5.7 (range 4.0 – 8.0) species compared to 4.6 (range 2.0 – 8.0) in fish-bearing lakes. Pielou’s evenness, Shannon-Weiner and Simpson’s indices showed no significant distinction between fishless and fish-bearing lakes.

Fishless and fish-bearing lakes supported distinct pelagic zooplankton communities (Fig. 4). Fish presence has a significant effect on zooplankton composition, measured by the Bray-Curtis dissimilarity (F_(1,33)_ = 0.798, p = 0.013). At the same time, connectivity did not show any effect (PERMANOVA, F_(1,33)_ = 0.316, p > 0.1).

**Figure 4.**
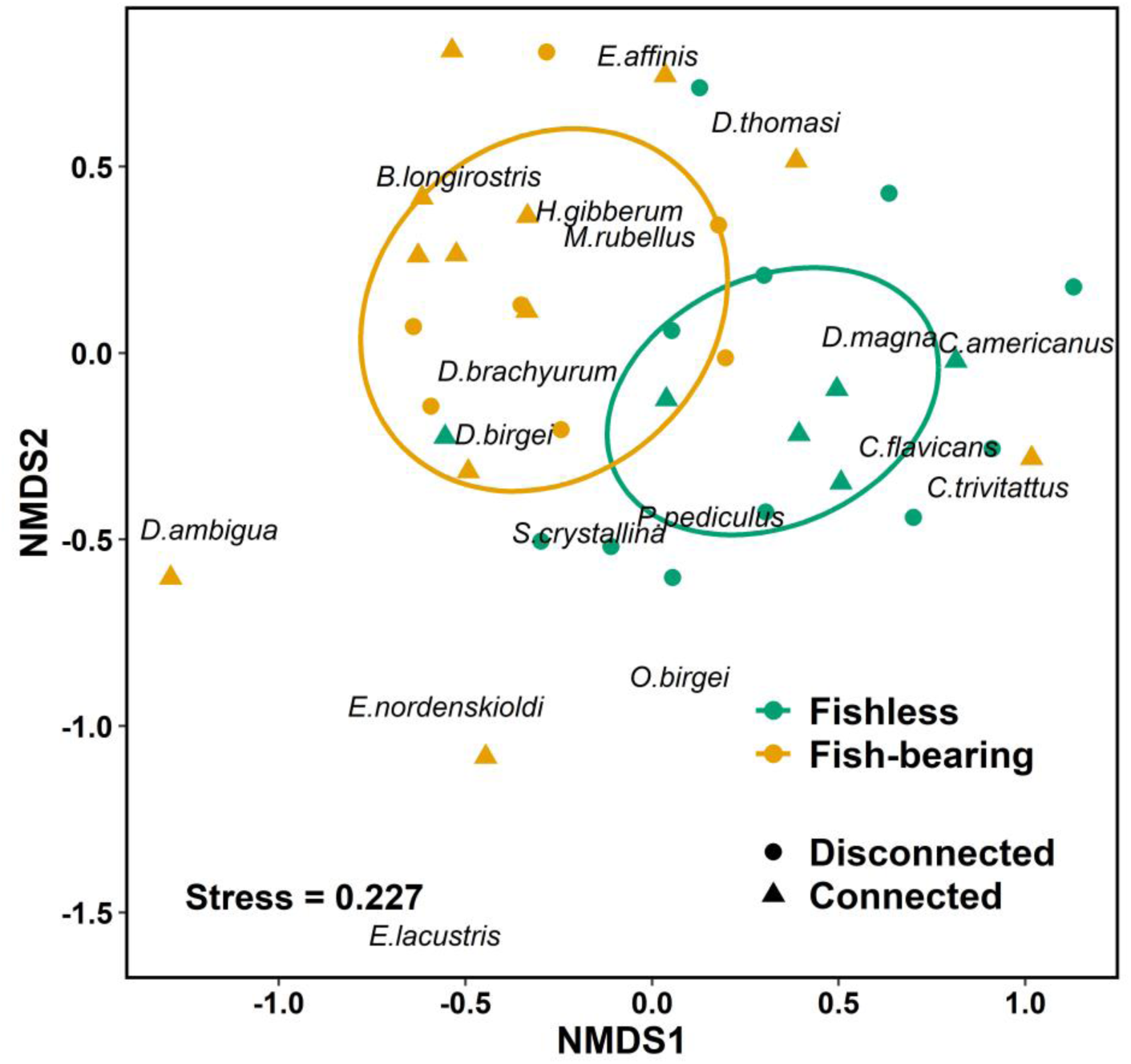
NMDS plot based on the Bray-Curtis dissimilarity metric for pelagic zooplankton communities in August 2024. Green: Fishless lakes (n=17), orange: Fish-bearing lakes (n=19), circle: Disconnected lakes (n=18), triangle: Connected lakes (n=18).

## Discussion

By comparing fishless and fish-bearing kettle lakes, we showed that the presence of fish impacts several key components of the lake food webs. Amphibians were affected at all life stages, probably because fish can either compete with them or prey on them at each stage. The egg and larval stages were the most critical, as fish prey on them more frequently than on adults. The presence of fish also altered zooplankton species composition, eradicated the most vulnerable species and reduced zooplankton diversity. Our conclusions confirmed our hypotheses and are supported by studies on fish introductions in high-elevation lakes (Bradford et al., 1998; Holmes et al., 2017; Knapp et al., 2001; Tiberti et al., 2014).

### Impacts of fish presence on amphibians at different life stages

We found that the fish presence in kettle lakes negatively affect amphibians throughout their life cycle. At their early life stages (eggs, hatching, larvae, and metamorphosis), amphibians are particularly exposed to direct predation by fish as they rely on aquatic environments (Hecnar & M’Closkey, 1997; Rodrigue & Desroches, 2018; Welsh Jr et al., 2006). In our study, the presence of fish was associated with lower amphibian egg mass and larval abundance, and with the absence of certain amphibian species at the larval stage. Specifically, egg masses of *Ambystoma* sp. were observed in both lake types, but larvae were absent in fish-bearing lakes. *Ambystoma* are palatable to fish (Kats et al., 1988), and fish predation is often sufficient to prevent larval recruitment (Maret et al., 2006; Pagnucco et al., 2011). Petranka (1983) demonstrated that fish can eliminate local population of smallmouth salamander larvae (*A. texanum*) in stream pool in the United States. Hecnar & M’Closkey (1997) have also shown that *Ambystoma* do not co-occur with fish. Adults may also be affected indirectly through competition and, at a lower level, directly by fish predation (Joseph et al., 2011; Welsh Jr et al., 2006).

Egg masses of the wood frog, a species palatable to fish (Hecnar & M’Closkey, 1997), were also found in lower abundance in fish-bearing lakes. A similar relationship between wood frog egg masses and fish was observed in ponds in the United States by Egan and Paton (2004). Mink frog tadpoles were also negatively impacted by fish, and their relative abundance decreased by 98% in fish-bearing lakes. Given its similarity to other boreal species studied here, we argue that fish also prey on mink frog tadpoles, although no studies have directly examined this. In light of these results, we can conclude that fish presence negatively impacts amphibian reproduction, most likely through the consumption of eggs and larvae, strongly compromising the reproductive success of amphibians in kettle lakes.

In addition to direct predation on eggs and larvae, adult amphibians could also avoid breeding in fish-bearing lakes. For instance, spotted salamanders deposit more egg masses in fishless ponds, and wood frogs preferentially select fishless breeding sites (Egan et Paton, 2004; Hopey et Petranka, 1994). Adult wood frogs can detect the presence of fish, likely via chemical or mechanical cues, and choose fishless habitats to maximize the fitness of their offspring (Hopey & Petranka, 1994). However, despite the negative impact of fish on amphibian eggs and larvae, we found that adult amphibians call in lakes independently of the presence or absence of fish. Adults are less susceptible to predation than eggs and larvae, as they are larger in size and more mobile (Hecnar & M’Closkey, 1997; Kloskowski, 2011; Welsh Jr et al., 2006), which may explain why they can reproduce in both lake types. Nevertheless, in our study, the effect of fish presence on the abundance of adult male anurans was species-dependent.

Three hypotheses can explain why the relationship between adult anuran abundance and fish presence is species-dependent. First, it may result from complex interactions among amphibian species (predation and competition), leading to perceived indirect effects of fish presence. For example, to avoid predation of their eggs and larvae by wood frogs, American toads select breeding sites where wood frogs are absent (Petranka et al., 1994). To avoid fish predation, wood frogs preferentially select fishless breeding sites (Hopey & Petranka, 1994). Consequently, American toad abundance could be indirectly and positively influenced by the presence of fish, as observed in our study. Other predatory interactions among amphibians species are well documented (Rodrigue & Desroches, 2018; Werner & McPeek, 1994), as well as interspecific competition for resources (Arribas et al., 2015). Second, amphibian species differ in palatability to fish (Hecnar & M’Closkey, 1997), which could lead to variable responses of adult male anurans to the presence of fish. However, our results do not fully support this hypothesis because green frogs, which are unpalatable to fish (Kats et al., 1988), were negatively affected by their presence, while northern spring peepers, which are palatable (Kats et al., 1988), were not affected. This is supported by Feldman et al. (2023), who did not detect a relationship in beaver and peatlands ponds. Finally, amphibian species may differ in their antipredator behaviours, such as activity levels and escape responses, which can influence predator efficiency (Werner & McPeek, 1994).

Despite negative impacts on amphibian abundance throughout their life cycle, we did not find a significant effect of fish presence on amphibian species richness, even though we observed lower salamander egg mass abundance and no larvae in fish-bearing lakes. This result agrees with Babbitt et al. (2003), who found no significant effect of predatory fish on larval amphibian species richness in permanent wetlands, even though spotted salamanders and wood frogs occurred only in fishless wetlands. Amphibian species richness can remain unchanged, while the community composition changes.

Furthermore, Werner et al. (2007) argue that, because amphibians are vulnerable to fish, the presence of fish is a local factor contributing to species composition turnover through local extinctions. A lack of observed differences in species richness may also be due to the limited number of amphibian species present and the presence of small, opportunistic omnivorous fish in the study lakes. Supporting this hypothesis, Hecnar and M’Closkey (1997) showed that piscivorous fish reduced amphibian species richness, whereas non-piscivorous fish had no impact on richness. Furthermore, some lakes classified as fishless may have contained fish, which could influence our results, since fish sampling was conducted only once between May and July. However, this scenario is unlikely, as most lakes had been sampled for fish three years earlier, and legal stocking records were consulted.

### Impacts of fish presence on zooplankton

Fish can modify zooplankton communities, altering species composition, diversity, and abundance through top-down control. Visual foraging planktivorous fish and juveniles of piscivorous species prey on zooplankton, preferentially selecting large, mobile, and pigmented individuals (Bradford et al., 1998; Hylander et al., 2012). High-elevation lakes, similarly to kettle lakes on esker, are often oligotrophic with clear water, forcing organisms to protect themselves from ultraviolet radiations through the accumulation of pigments such as carotenoids or melanin (Sommaruga, 2010; Ulbing et al., 2019) which make them more conspicuous and vulnerable to fish predation (Grosbois & Rautio, 2018; Schneider et al., 2016, 2017). Consequently, the introduction of fish into naturally fishless lakes may exert a strong predatory pressure on zooplankton such as *Leptodora* and phantom midges (*Chaoborus* larvae), as well as large-bodied zooplankton. Although we did not detect an impact on large-bodied Daphniidae in our system, we observed that *Chaoborus* species were less likely to occur in fish-bearing lakes, and *C. americanus* was captured only in fishless lakes. This species is highly vulnerable to fish predation because it cannot escape through diel vertical migration in the water column (von Ende, 1979). Fish predation therefore regulates both the abundance and distribution of *C. americanus* (Drouin et al., 2009). Holmes et al. (2017), Drouin et al. (2009) and Schilling et al. (2009b) also reported that *C. americanus* was absent from most fish-bearing lakes. Consequently, the reduced predation pressure by *Chaoborus* on Daphniidae in fish-bearing lakes, combined with direct fish predation on Daphniidae, may explain the absence of a pattern in Daphniidae abundance among lake types. This result contrasts with Drouin et al. (2009), who found that *Chaoborus* predation had a greater impact on large-bodied zooplankton than did fish predation in Eastern Boreal Shield lakes. Drouin et al. (2009) attributed this difference to the efficiency and intensity of fish predation on zooplankton. In our study, opportunistic omnivorous fish likely explain the strong impact of fish presence on zooplankton community structure.

As fish prey on *Chaoborus* larvae, and thus reduce predation pressure by *Chaoborus* on zooplankton, we found a higher abundance of small-bodied zooplankton, such as Bosminidae and Diaptomidae in fish-bearing lakes. This pattern is well documented in the literature, where fish reduce the abundance of large-bodied zooplankton and, thus, enhance the abundance of smaller-bodied zooplankton (J. L. Brooks & Dodson, 1965; Tiberti et al., 2014). Through their top-down control, fish presence also influences zooplankton species composition, which differs between fishless and fish-bearing lakes. *Chaoborus* species and *Daphnia magna* seem to have a strong affinity for fishless lakes and cladocerans, as *Diaphanosoma* sp., *Holopedium gibberum* and Bosminidae, were more characteristic of fish-bearing lakes, which agreed with Mushet et al. (2020).

Zooplankton species richness was also reduced by the presence of fish in kettle lakes. As opposed to other experiments that noticed an increase in fish-bearing lakes or no difference in zooplankton diversity between fishless and fish-bearing lakes (Drouin et al., 2009; Holmes et al., 2017), kettle lakes on esker have very low zooplankton abundance (Couture et al., 2021; Drouin et al., 2009). We detected as few as 78 individuals per 1 000 L in some lakes. It is possible that, through their introduction and the new predation pressure they exert, fish eradicate the most vulnerable zooplankton species or the least abundant ones. Moreover, in our study, Pielou’s evenness, Shannon-Weiner and Simpson’s indices are similar between lakes with and without fish. These evenness and diversity indices are not sensitive to rare taxa, which can be easily eliminated by fish predation due to their low abundance. Yet, the presence of fish may have reduced the abundance or eliminated rare species without being reflected in the evenness and diversity indices.

### Impacts of connectivity on amphibians and zooplankton

Connectivity was expected to modulate the impacts of fish on amphibian and zooplankton communities by allowing continuous movement of individuals. However, our results did not support this hypothesis, as connectivity had no significant effect on our communities, except for adults’ American toads, adults’ wood frogs and mink frog larvae.

Werner et al. (2007) showed that pond connectivity was positively correlated with turnover in amphibian species composition, as it facilitates amphibian dispersal and colonization of new ponds. The lack of a significant effect of connectivity in our study, as well as its interaction with fish presence, for most amphibian species may indicate that the strong negative impact of fish on amphibian abundance has outweighed any potential positive effect of connectivity, making the latter undetectable. It is also possible that the connectivity, as we measured it, did not reflect the actual dispersal corridors used by amphibian species, as some species preferentially use terrestrial pathways (Rodrigue & Desroches, 2018). However, the significant positive interaction between fish and connectivity for mink frog larvae and calling males of American toad and wood frog suggests that the impact of fish differs between naturally colonized (connected lakes) and stocked (disconnected) lakes. Amphibian communities may be more adapted to long-established, naturally occurring fish populations than to recently introduced stocked fish (Závorka et al., 2018), which could explain why we observed a weaker impact of fish in connected lakes. Historical records of fish presence in these lakes would have provided valuable insight to confirm this hypothesis and to better understand the long-term versus short-term impacts of fish presence on aquatic communities.

Zooplankton have limited dispersal capacities (Havel & Shurin, 2004), constraining their ability to colonize new waterbodies. Thus, no effect of connectivity, or of its interaction with fish presence, on zooplankton abundance, diversity, and community composition was detected in our study.

### Impacts of environmental variables on amphibians and zooplankton

We showed that DOC concentration was positively associated with relative larval abundance of amphibians. DOC is known to hinder light penetration in water and to attenuate ultraviolet B radiation (UVBR) in the water column (Croteau et al., 2008). Consequently, it may release the predation pressure of visual predators on amphibian larvae in kettle lakes on esker that have clear water. As high DOC concentrations degrade lake optical conditions, it reduces foraging success by fish (Weidel et al. (2017), enhancing survivorship of amphibian larvae. Moreover, as UVBR exposure reduced survivorship of amphibian larvae (Croteau et al., 2008), an increase in DOC can positively impact their relative larval abundance.

Other measured physico-chemical parameters did not impact the amphibian community in our study. However, although both the data and the literature support our hypotheses, the apparent effect of fish presence on amphibians may partly reflect the influence of other unmeasured environmental variables. For instance, Egan and Paton (2004) showed that vegetation cover positively influenced oviposition of wood frogs. Kettle lakes on esker are typically characterized by low macrophyte density with low variability (Hasan et al., 2023), suggesting that vegetation cover likely plays a minor role in determining egg mass distribution in these ecosystems. Lake size and depth could also influence amphibian communities by providing more habitats and supporting higher population densities (Laan & Verboom, 1990). However, Knapp et al. (2003) reported that water depths greater than 4 m have little influence on amphibians. The kettle lakes in our study were almost all deeper than this threshold (AB, personal observation). To minimize the potential confounding effect of lake size, we also selected similar kettle lakes, all smaller than 7 ha. Finally, the shoreline slope may affect amphibian dispersal, as steep shorelines can act as physical barriers by increasing the energetic cost of movement between aquatic and terrestrial environments (Lowe et al., 2006). Because some of the studied kettle lakes exhibited very steep slopes while other had much gentler slopes, slope may have influenced amphibian community structure.

### Conclusions

Our study demonstrated that fish presence in kettle lakes affected amphibian and zooplankton communities, and this pattern was not modulated by hydrological connectivity. Amphibians were impacted by the presence of fish throughout their life cycle. Adult amphibians can reproduce in both fish-bearing and fishless lakes, but the lower number of egg masses and the reduced relative larval abundance observed in fish-bearing lakes suggest that fish presence may compromise reproductive success and can threaten the long-term resilience of amphibian communities in kettle lakes. Fishless kettle lakes can therefore constitute critical habitats for amphibians, particularly for salamander larvae, which were absent from fish-bearing lakes. Similarly, *Chaoborus americanus*, a predatory zooplankton species, occurred exclusively in fishless kettle lakes, indicating that these lakes also support unique components of the zooplankton community. Because naturally fishless kettle lakes are unique ecosystems in the landscape, we thus recommend avoiding fish stocking in these lakes to preserve essential reproductive habitats for amphibians, maintain refuges for sensitive zooplankton species, and safeguard the spatial heterogeneity that underpins landscape-scale biodiversity.

## Acknowledgements

We thank all the interns, students and employees from GREMA who made this research possible: Audrey Beaudette, Amé Bergeron, Élise Berthiaume, Lehann Bouchard, Noé Bruel, Louis-Philippe Charest, Thomas Dubé, Martine Hardy, Julie-Pascale Labrecque-Foy, Gabriela Soucy-Cardoso, Myriam Marentette, Marie-Claude Mayotte and Mathias Mayen. Fish and amphibians sampling protocols were approved in advance by the ethics committee of UQAT and two sampling permits were obtained from the Québec Ministère de l’Environnement, de la Lutte contre les changements climatiques, de la Faune et des Parcs (2024-04-17-024-08-GP; 2024-04-17-028-08-GF).

## Author Contributions

Conceptualisation: AB, GG. Developing methods and writing: AB, GG, KT, MF. Conducting the research, data analysis and preparation of figures and tables: AB. Data interpretation: AB, GG, KT.

## Data availability statement

Data are available from the corresponding author upon reasonable request.

