## Supplementary material for "Fish presence alters amphibian and zooplankton communities in kettle lakes, but not hydrological connectivity"

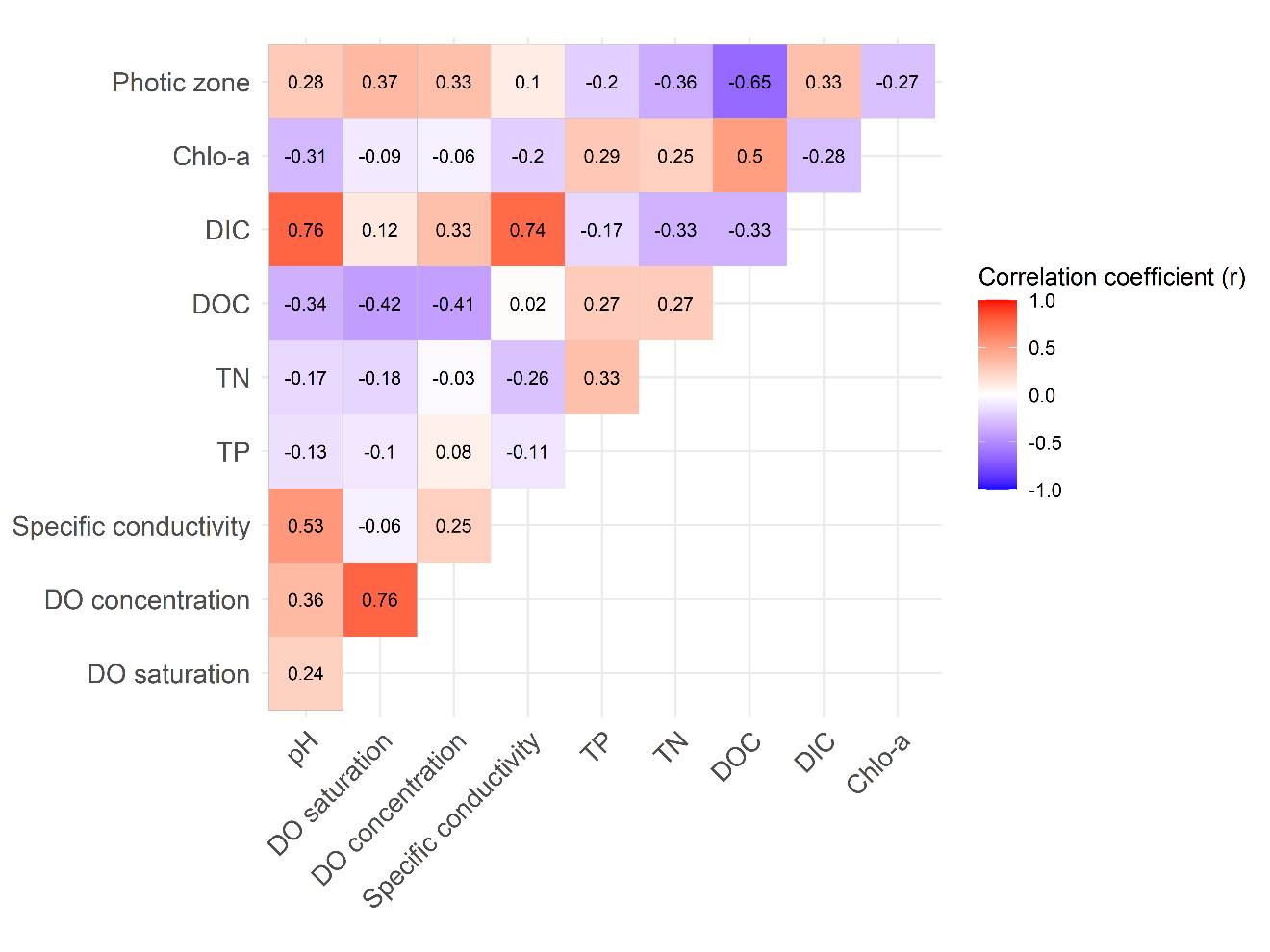

**Figure S1.** Pearson correlation matrix of physico-chemical variables. Variables were considered correlated if |r| > 0.7, where *r* is the correlation coefficient. Abbreviations: Chlo-a, chlorophyll-a; DIC, dissolved inorganic carbon; DOC, dissolved organic carbon; TN, total nitrogen; TP, total phosphorus; DO, dissolved oxygen.

**Table S1.** Selection of independent variables to be included in statistical models based on biological relevance of the variables (Bevier et al., 2022; Dodd, 2023; Mills & Ward, 2015; Roby-Thomas, 2006), Boruta feature selection algorithm, the best model derived from Akaike Information Criterion (AIC), and backward elimination. * indicates and interaction between the two variables.

| **Dependant variables** | **Independent variables to be included in statistical models based on:** | | | |
| --- | --- | --- | --- | --- |
|  | **Biological relevance of the variables** | **Boruta feature selection algorithm** | **Best model based on Akaike Information Criterion (AIC) values** | **Backward elimination** |
| **Egg masses’ presence** | Fish presence | Dissolved oxygen concentration | Fish presence | Fish presence |
|  | Connectivity | Specific conductivity | Dissolved oxygen concentration | Connectivity |
|  | Fish presence*connectivity | Dissolved inorganic carbon | pH | Species |
|  | Species |  |  |  |
|  | Dissolved oxygen concentration |  |  |  |
|  | Specific conductivity |  |  |  |
|  | Dissolved organic carbon  pH |  |  |  |
| **Spotted salamander egg masses’ abundance** | Fish presence | Fish presence | Fish presence | Fish presence |
|  | Connectivity | Specific conductivity | Connectivity |  |
|  | Fish presence*connectivity | Dissolved oxygen saturation | Fish presence * connectivity |  |
|  | Dissolved oxygen concentration | Dissolved oxygen concentration | pH |  |
|  | Dissolved organic carbon | Dissolved inorganic carbon |  |  |
|  | pH |  |  |  |
| **Wood frog egg masses’ abundance** | Fish presence |  | Fish presence | Fish presence |
|  | Connectivity |  | Specific conductivity | Connectivity |
|  | Fish presence*connectivity |  | pH |  |
|  | Dissolved oxygen concentration |  |  |  |
|  | Specific conductivity |  |  |  |
|  | pH |  |  |  |
| **Mink frog egg masses’ abundance** | Fish presence |  | Fish presence | Fish presence |
|  | Connectivity |  | Connectivity | Connectivity |
|  | Fish presence * connectivity |  | Fish presence*connectivity |  |
|  | Specific conductivity |  | Specific conductivity |  |
|  | pH |  | pH |  |
| **Mink frog tadpole relative abundance** | Fish presence | Fish presence | Fish presence | Fish presence |
|  | Connectivity |  | pH | Connectivity |
|  | Fish presence * connectivity  Specific conductivity  pH |  |  | Fish presence*connectivity |
| **Green frog tadpole relative abundance** | Fish presence | Dissolved oxygen concentration | Dissolved organic carbon | Fish presence |
|  | Connectivity | Photic zone |  | Connectivity |
|  | Fish presence*connectivity |  |  |  |
|  | Dissolved organic carbon |  |  |  |
|  | pH |  |  |  |
| **Amphibian relative larval abundance** | Fish presence | Fish presence | Fish presence | Fish presence |
|  | Connectivity | Dissolved organic carbon | Dissolved organic carbon | Dissolved organic carbon |
|  | Fish presence *connectivity | Specific conductivity |  |  |
|  | Dissolved oxygen concentration |  |  |  |
|  | Specific conductivity  Dissolved organic carbon  pH |  |  |  |

**Table S2.** Relative abundance (catch per 24-hour period) of fish species and mean species richness in connected and disconnected lakes (mean ± standard deviation [min, max]).

| Species | Lake type | |
| --- | --- | --- |
|  | Connected lakes (n=12) | Disconnected lakes (n=7) |
| Brook stickleback  (*Culaea inconstans*) | 84.8 ± 152.6 [0, 462] | 247.4 ± 391.0 [0, 1053] |
| Brown bullhead  (*Ameiurus nebulosus*) | 35.0 ± 94.5 [0, 322] | 114.3 ± 229.8 [0, 610] |
| Northern redbelly dace (*Chrosomus eos*) | 58.3 ± 186.3 [0, 649] | 14.6 ± 27.7 [0, 72] |
| Yellow perch  (*Perca flavescens*) | 73.7 ± 125.1 [0, 424] | 27.4 ± 52.2 [0, 136] |
| White sucker  (*Catostomus commersoni*) | 10.5 ± 36.4 [0, 126] | 8.0 ± 19.5 [0, 52] |
| Pearl dace  (*Margariscus margarita*) | 58.8 ± 153.6 [0, 514] | 0 |
| Golden shiner  (*Notemigonus crysoleucas*) | 16.9 ± 58.6 [0, 203] | 0 |
| Mottled sculpin  (*Cottus bairdi*) | 0.2 ± 0.6 [0, 2] | 0 |
| Fathead Minnow  (*Pimephales promelas*) | 1.3 ± 4.6 [0, 16] | 0 |
| Brook trout  (*Salvelinus fontinalis*) | 1.2 ± 4.0 [0, 14] | 0 |
| Others Cyprinidae | 0 | 4.0 ± 10.6 [0, 28] |
| Species richness | 2.2 ± 1.3 [1, 5] | 1.7 ± 1.3 [1, 4] |

**Table S3** Models results showing the independent variables included in the most parsimonious model for each dependent variable, estimate values and their standard error (β ± SE), and associated p-values. An Akaike Information Criterion (AIC) value was obtained for every fitted model. Akaike weights (ω_i_) were calculated only for the models in which all possible variable combinations were compared and ranked by AIC to perform variable selection. For wood frog egg masses’ abundance and green frog tadpole relative abundance models, Akaike weights wasn’t calculated as too many models didn’t converge. Statistically significant variables are notes in bold. *: ≤ 0.05; **: p ≤ 0.01; ***: p ≤ 0.001.

| **Dependent variables** | **Independent variables** | ***β* ± SE** | ***p*-values** | **AIC** | **ω_i_** |
| --- | --- | --- | --- | --- | --- |
| **Amphibians** | | | | | |
| **Zero-inflated negative binomial model** | | |  | |  |
| Spotted salamander egg masses’ abundance | Fish presence | -2.55 ± 1.15 | **0.027*** | 143.0 | 0.002 |
| Wood frog egg masses’ abundance | Fish presence  Connectivity | -3.95 ± 2.11  1.75 ± 2.11 | 0.06  0.41 | 110.2 | *NA* |
| Mink frog egg masses’ abundance | Fish presence  Connectivity | -0.14 ± 0.61  -0.10 ± 0.59 | 0.82  0.87 | 68.4 | 0.003 |
| Mink frog tadpole relative abundance | Fish presence  Connectivity  Fish presence*connectivity | -3.72 ± 1.06  0.22 ± 0.79  2.95 ± 1.38 | **<0.001*****  0.775  **0.032*** | 137.0 | 0.014 |
| Green frog tadpole relative abundance | Fish presence  Connectivity | -1.72 ± 4.43  2.39 ± 4.42 | 0.589  0.698 | 82.5 | *NA* |
| **GLM, binomial family** | | |  | |  |
| Egg masses’ presence | Fish presence  Connectivity  Species  Mink frog  Wood frog | -1.23 ± 0.50  0.46 ± 0.49  -0.35 ± 0.59  0.30 ± 0.55 | **0.014***  0.342  0.550  0.587 | 117.8 | <0.001 |
| **LM** *(p of the model: <0.001)* | | |  | |  |
| log (amphibian relative larval abundance + 1) | Fish presence  Dissolved organic carbon | -2.11 ± 0.34  0.51 ± 0.17 | **<0.001*****  **0.006**** | 97.2 | 0.05 |
| **GAM, OCAT family** | | |  | |  |
| Northern spring pepper calling score (1^st^ sampling period) | **Linear predictors**  Fish presence  Connectivity  Fish presence*connectivity  **Non-linear predictor**  Julian day | 0.35 ± 0.80  -0.97 ± 0.81  -0.56 ± 1.18 | 0.658  0.230  0.635  **<0.001***** | 114.9 |  |
| Northern spring pepper calling score (2^nd^ sampling period) | **Linear predictors**  Fish presence  Connectivity  Fish presence *connectivity  **Non-linear predictor**  Julian day | -0.50 ± 0.38  -0.13 ± 0.40  0.38 ± 0.56 | 0.182  0.747  0.494  **<0.001***** | 420.7 |  |
| Wood frog calling score | **Linear predictors**  Fish presence  Connectivity  Fish presence *connectivity  **Non-linear predictor**  Julian day | -1.35 ± 0.37  -0.37 ± 0.36  1.16 ± 0.53 | **<0.001*****  0.305  **0.03***  **<0.001***** | 547.7 |  |
| American toad calling score (1^st^ sampling period) | **Linear predictors**  Fish presence  Connectivity  Fish presence *connectivity  **Non-linear predictor**  Julian day | 2.04 ± 0.65  1.58 ± 0.68  -0.98 ± 0.82 | **0.002****  **0.019***  0.232  **<0.001***** | 250.7 |  |
| American toad calling score (2^nd^ sampling period) | **Linear predictors**  Fish presence  Connectivity  Fish presence *connectivity  **Non-linear predictor**  Julian day | -1.66 ± 0.69  -0.54 ± 0.60  3.19 ± 0.94 | **0.016***  0.368  **<0.001*****  **<0.001***** | 207.6 |  |
| Mink frog calling score | **Linear predictors**  Fish presence  Connectivity  Fish presence *connectivity  **Non-linear predictor**  Julian day | 0.72 ± 0.36  0.18 ± 0.39  0.02 ± 0.53 | **0.045***  0.647  0.969  **<0.001***** | 544.0 |  |
| Green frog calling score | **Linear predictors**  Fish presence  Connectivity  Fish presence *connectivity  **Non-linear predictor**  Julian day | -1.51 ± 0.55  -0.21 ± 0.44  1.31 ± 0.73 | **0.006****  0.640  0.073  0.126 | 314.3 |  |
| **Zooplankton** | | | | |  |
| **GLM, binomial family** | | |  | |  |
| *Chaoborus* larvae presence | Fish presence | -2.20 ± 0.77 | **0.005**** | 44.2 |  |
| **LM** *(p of the model: 0.037)* | | |  | |  |
| log (species richness + 1) | Fish presence | -0.22 ± 0.10 | **0.037*** | 21.3 |  |

**Table S4.** Relative abundance (catch per 1000 L of filtered water) of zooplankton taxa in connected and disconnected, fish-bearing and fishless lakes (mean ± standard deviation [min, max]).

| Taxa | Lake type | | | |
| --- | --- | --- | --- | --- |
|  | Fishless and disconnected lakes (n=11) | Fishless and connected lakes (n=6) | Fish-bearing and disconnected lakes (n=7) | Fish-bearing and connected lakes (n=12) |
| Female calanoids | 274.9 ± 461.8 [0, 1 225] | 1032.2 ± 1 828.8 [0, 4 598] | 1 253.6 ± 960.7 [0, 2 570] | 240.0 ± 433.4 [0, 1 178] |
| Male cyclopoids | 21.5 ± 42.7 [0, 143] | 0.5 ± 0.8 [0, 2] | 60.1 ± 66.3 [0, 184] | 3.8 ± 8.7 [0, 29] |
| Nauplii | 589.6 ± 958.4 [10, 3 340] | 929.8 ± 763.7 [97, 1 678] | 1 892.1 ± 2 047.2 [51, 5 543] | 1 136.8 ± 1 879.9 [11, 5 386] |
| Juvenile calanoids | 146.2 ± 280.5 [0, 924] | 318.0 ± 647.0 [0, 1 630] | 478.3 ± 794.5 [0, 2 256] | 55.9 ± 90.5 [0, 315] |
| Juvenile cyclopoids | 94.5 ± 119.2 [7, 399] | 113.5 ± 203.1 [2, 522] | 304.3 ± 393.5 [0, 1 126] | 1 114.4 ± 3 580.5 [0, 12 474] |
| Cladocerean sp. | 0 | 0 | 0.1 ± 0.4 [0, 1] | 3.0 ± 10.4 [0, 36] |
| Sididae sp. | 7.8 ± 25.9 [0, 86] | 0 | 0.9 ± 2.3 [0, 6] | 0.2 ± 0.9 [0, 3] |
| Bosminidae sp. | 1.6 ± 3.5 [0, 11] | 0 | 0.7 ± 1.9 [0, 5] | 119.8 ± 415.1 [0, 1 438] |
| Chydoridae sp. | 13.9 ± 33.3 [0, 104] | 1.0 ± 2.5 [0, 6] | 3.7 ± 5.8 [0, 14] | 0 |
| Temoridae sp. | 0 | 0 | 9.6 ± 25.3 [0, 67] | 0.5 ± 1.7 [0, 6] |
| *Diaphanosoma birgei* | 21.9 ± 43.5 [0, 142] | 9.7 ± 18.4 [0, 46] | 129.6 ± 268.5 [0, 732] | 2.8 ± 6.0 [0, 21] |
| *Holopedium gibberum* | 5.8 ± 5.1 [0, 16] | 113.0 ± 244.1 [0, 610] | 793.0 ± 1 259.3 [0, 2 922] | 188.3 ± 347.2 [0, 1 124] |
| *Polyphemus pediculus* | 90.6 ± 295.4 [0, 981] | 1.7 ± 3.2 [0, 8] | 0 | 5.8 ± 13.4 [0, 35] |
| *Ceriodaphnia* sp. | 28.9 ± 83.2 [0, 278] | 3.2 ± 7.8 [0, 19] | 67.9 ± 176.9 [0, 469] | 83.6 ± 228.2 [0, 799] |
| *Daphniidae* sp. | 0 | 0 | 1.3 ± 3.4 [0, 9] | 0 |
| *Daphnia ambigua* | 0 | 0.3 ± 0.8 [0, 2] | 5.1 ± 11.9 [0, 32] | 136.6 ± 378.7 [0, 1 298] |
| *Daphnia magna* | 13.0 ± 27.6 [0, 94] | 1.0 ± 1.6 [0, 3] | 10.6 ± 26.2 [0, 70] | 2.9 ± 10.1 [0, 35] |
| *Cyclopidae* sp. | 52.9 ± 122.7 [0, 411] | 12.2 ± 15.6 [0, 38] | 165.1 ± 214.9 [0, 604] | 20.4 ± 38.0 [0, 107] |
| *Diacyclops thomasi* | 9.7 ± 26.3 [0, 88] | 0.7 ± 1.2 [0, 3] | 21.0 ± 39.2 [0, 101] | 21.1 ± 72.4 [0, 251] |
| *Microcyclops rubellus* | 9.1 ± 22.5 [0, 72] | 1.5 ± 2.5 [0, 6] | 8.9 ± 21.0 [0, 56] | 3.8 ± 9.0 [0, 27] |
| *Microcyclops* sp*.* | 0 | 0 | 24.4 ± 50.9 [0, 136] | 0 |
| *Chaoborus flavicans* | 10.0 ± 19.7 [0, 67] | 33.2 ± 30.3 [0, 74] | 1.1 ± 2.0 [0, 5] | 2.8 ± 9.5 [0, 33] |
| *Chaoborus americanus* | 2.6 ± 6.9 [0, 23] | 0.3 ± 0.8 [0, 2] | 0 | 0 |
| *Chaoborus* sp. | 4.4 ± 9.5 [0, 30] | 4.0 ± 5.6 [0, 15] | 1.0 ± 1.3 [0, 3] | 1.2 ± 4.0 [0, 14] |
| *Chaoborus trivitattus* | 4.3 ± 11.2 [0, 37] | 6.2 ± 9.9 [0, 26] | 0 | 1.7 ± 5.8 [0, 20] |
| *Bosmina longirostris* | 86.7 ± 130.9 [0, 20 340] | 250.7 ± 525.1 [0, 1 319] | 1230.7 ± 1 403.6 [14, 3 235] | 1 988.5 ± 5 922.9 [0, 20 773] |
| *Epischura nordenskioldi* | 0 | 0 | 4.6 ± 12.1 [0, 32] | 8.1 ± 20.8 [0, 69] |
| *Epischura lacustris* | 0 | 0 | 0 | 0.4 ± 1.4 [0, 5] |
| *Diaptomidae* | 19.7 ± 49.8 [0, 162] | 43.2 ± 67.0 [0, 172] | 223.7 ± 322.0 [0, 898] | 26.5 ± 46.1 [0, 133] |
| *Sida crystallina* | 6.6 ± 14.7 [0, 40] | 0 | 0 | 0.9 ± 3.2 [0, 11] |
| *Onychodiaptomus birgei* | 0.1 ± 0.3 [0, 1] | 0 | 0 | 0 |
| *Eurytemora affinis* | 0.1 ± 0.3 [0, 1] | 0 | 1.7 ± 3.4 [0, 9] | 0.7 ± 2.3 [0, 8] |
